# Phyloecology of *nrfA*-ammonifiers and their relative importance with denitrifiers in global terrestrial biomes

**DOI:** 10.1101/2023.04.08.536100

**Authors:** Aurélien Saghaï, Grace Pold, Christopher M. Jones, Sara Hallin

**Affiliations:** Swedish University of Agricultural Sciences, Department of Forest Mycology and Plant Pathology, Uppsala, Sweden

**Keywords:** phylogeny, metagenome, *nrfA*, soil, machine learning, DNRA

## Abstract

Nitrate ammonification is important for soil nitrogen retention. However, the ecology of nitrate ammonifiers and their prevalence compared with denitrifiers, being competitors for nitrate, are overlooked. Here, we screened more than 1 million genomes for *nrfA*, encoding the nitrite reductase in nitrate ammonification. Nearly 50% of the nitrate ammonifier assemblies carry at least one denitrification gene and, contrary to the current paradigm, have higher potential for nitrous oxide production than reduction. We then used a phylogeny-based approach to recruit *nrfA* and denitrification nitrite reductase gene fragments in 1,861 metagenomes covering the major terrestrial biomes. Denitrification genes dominated, except in tundra, and random forest modelling teased apart the influence of the soil C/N on nitrate ammonifier vs denitrifier abundances, showing an effect of nitrate rather than carbon content. This study demonstrates the multiple roles nitrate ammonifiers play in nitrogen cycling and the factors ultimately controlling the fate of nitrate in soil.

---

Human activity, in particular agricultural fertilizer application and fossil-fuel combustion, has increased the amount of nitrogen (N) circulating in the biosphere and created an imbalance in the N cycle that threatens ecosystem integrity at the global scale (Steffen *et al*., 2015). Nitrate is a highly mobile form of reactive N in soil and the primary source of global N pollution (Kanter *et al*., 2020). If not assimilated into biomass or leached to watersheds, nitrate can be used as an electron acceptor by soil microorganisms under oxygen-limited conditions, mainly through denitrification or nitrate ammonification, also known as dissimilatory nitrate reduction to ammonium. Denitrification leads to N loss through the production of gaseous N-compounds, including the potent greenhouse gas nitrous oxide (N2O). Terrestrial ecosystems contribute ca. 60 % to global N_2_O emissions (Tian *et al*., 2020), with denitrification being the main source, and N_2_O concentration in the atmosphere is increasing at an accelerating rate (Thompson *et al*., 2019). By contrast, nitrate ammonification does not generate intermediates and results in the retention of N via the binding of ammonium to negatively charged surfaces in the soil. Determining the environmental factors controlling the end-products when nitrate is used as electron acceptor is crucial for our ability to predict and influence N budgets in terrestrial ecosystems at the global scale (Battye *et al*., 2017). A key factor is a better understanding of the ecology of nitrate-ammonifying microorganisms, as they are an overlooked functional group in the N cycle (Kuypers *et al*., 2018), in particular in terrestrial ecosystems.

Dissimilatory nitrate ammonification is primarily driven by microorganisms using the periplasmic pentaheme cytochrome c nitrite reductase NrfA, encoded by the *nrfA* gene, to catalyze the reduction of nitrite to ammonium (Simon & Kroneck, 2014). Similar to denitrification, nitrate is reduced to nitrite by reductases encoded by *narG* or *napA*, and the branching point between the two pathways is the reduction of nitrite. While nitrate ammonification involves the reduction of nitrite to ammonium in a single enzymatic step, denitrification is a modular pathway in which nitrite is successively reduced into nitric oxide, N_2_O and finally dinitrogen gas via reductases encoded by *nirK* or *nirS*, *nor* and *nosZ*, respectively (Graf *et al*., 2014). Both nitrate ammonification driven by NrfA and the competing process denitrification are performed by phylogenetically diverse bacteria and archaea (Philippot *et al*., 2007; Welsh *et al*., 2014). NrfA-driven ammonification has been suggested to dominate or increase in relation to denitrification under electron acceptor limitation, i.e. at high ratios of soil organic carbon (SOC) and nitrate, whereas conditions with electron donor limitation favor denitrification (Tiedje *et al*., 1982). This is supported by more recent work with enrichment and pure cultures (Stremińska *et al*., 2012; Kraft *et al*., 2014; van den Berg *et al*., 2015; Yoon *et al*., 2015), site-specific field studies (Putz *et al*., 2018; Pandey *et al*., 2019; Luo *et al*., 2020) and modeling approaches (Algar & Vallino, 2014; Jia *et al*., 2020). Yet, we lack a synthesis of the relative importance of the SOC to nitrate ratio and other factors for the competition to use nitrate as an electron acceptor across Earth’s terrestrial biomes (Yang *et al*., 2017).

In this study, we determine the extant diversity, abundance, and global distribution of *nrfA*-ammonifiers and the environmental drivers of the potential competition with denitrifiers in terrestrial ecosystems. This includes an extensive phylogenetic analysis of full-length *nrfA* sequences obtained from screening more than 1,000,000 assemblies of isolate and metagenome-assembled genomes (MAGs). Both pathways have been shown to co-exist in a few but phylogenetically diverse isolates (Sanford *et al*., 2012; Yoon *et al*., 2015; Mania *et al*., 2016). Therefore, the presence of denitrification genes in the assemblies was determined to gain insights into the overall patterns of N_2_O production and reduction capacity among *nrfA*-ammonifiers. In addition to the *nrfA* phylogeny, we also use updated phylogenies of the genes *nirS* and *nirK* (Graf *et al*., 2014), coding for the equivalent function in denitrifiers, to provide a phylogenetic framework for analyzing ammonifying and denitrifying microorganisms. This approach circumvents the use of annotation pipelines with arbitrary cutoff values that may have little biological significance in metagenomic studies (e.g. Nelson *et al*., 2016). The phylogenies were used as references for recruiting bacterial and archaeal *nrfA, nirS*, and *nirK* fragments from 1,861 soil and rhizosphere metagenomes derived from broad environmental gradients across 725 locations to assess the global distribution of *nrfA*-ammonifiers and their abundance relative to denitrifying microorganisms. We further identify environmental drivers underpinning the abundance of functional microbial communities performing NrfA-driven ammonification *vs.* denitrification as a proxy for the competition between these groups using random forest modelling and discuss the implications of the findings for N loss and retention in global soils.

## Results

### Phylogeny of NrfA

The search for the presence of *nrfA* in isolate genomes and MAGs resulted in 1,150 non-redundant taxonomically and structurally diverse sequences from 1,121 assemblies, spanning 1 archaeal and 43 bacterial phyla (**Extended Data Table 1**). Only 24 assemblies (ca. 2 %) carried more than one copy of *nrfA*, with high sequence similarity between copies (**Extended Data Fig. 1**). The NrfA phylogeny was overall consistent with that of the organisms at the class level, except for some taxa including Anaerolineae, Campylobacteria, Gammaproteobacteria and Myxococcia (**Fig. 1**). While all sequences contained the five heme-binding sites and a histidine residue between the third and fourth site that are characteristic of NrfA (Einsle *et al*., 1999), a number of other structural features were associated with different clades within the NrfA phylogeny. Sequences with a Cys-X-X-Cys-His (CXXCH) motif in the first site, instead of the more common Cys-X-X-Cys-Lys, were exclusively bacterial and formed a monophyletic, well supported and taxonomically diverse clade (**Fig. 1**) (Welsh *et al*., 2014; Soares *et al*., 2022). Most known NrfA proteins are characterized by the presence of a calcium ion near the active site, where it is suspected to play a structural role (Cunha *et al*., 2003), whereas those that are calcium-independent contain a X-X-Arg-His motif between the third and fourth sites. The latter were present in four regions of the tree (n = 129 sequences), supporting independent evolutionary events (Campeciño *et al*., 2020).

**Figure 1.**
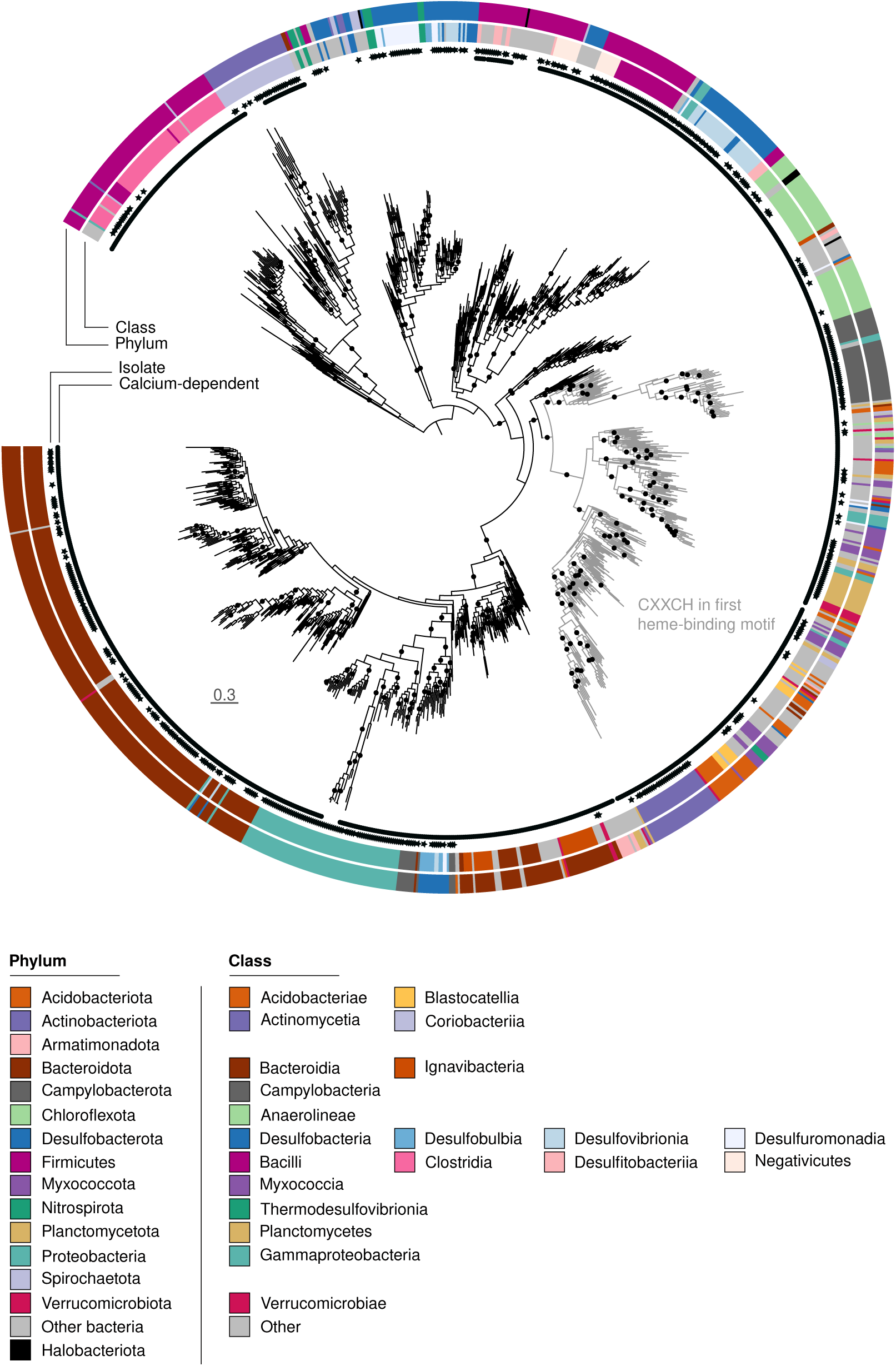
Maximum likelihood phylogeny of 1,150 full-length NrfA sequences from 1,121 genome assemblies inferred from the alignment of 512 amino acid positions. Calcium-dependent NrfA sequences are indicated by circles in the inner ring. Sequences obtained from isolates are shown by black stars and the rest are obtained from metagenome-assembled genomes. Taxonomic classification at the phylum and class level of the most abundant classes (n > 10, except for the archaeal class Halobacteriota) is indicated by the color in the two outer rings and is based on the Genome Taxonomy DataBase. Black circles on the phylogeny show support values (SH-aLRT test ≥ 80% and ultrafast bootstrap ≥ 95%, each threshold corresponding to an estimated confidence level of 95 %) and the scale bar denotes the amino acid exchange rate (WAG+R10). The outgroup is not shown.

### Potential for denitrification in genomes of nrfA-ammonifiers

Genomes harboring *nrfA* were further examined for the presence of denitrification genes. About 45 % of the assemblies harboring *nrfA* contained at least one denitrification gene (*nir*, *nor* or *nosZ*), whereas 13 % carried more than one denitrification gene, with complete denitrifiers accounting for just 2 % of the *nrfA*-ammonifiers (**Fig. 2a**). Microorganisms carrying both *nrfA* and *nir*/*nor* genes can contribute to either N retention or N loss in ecosystems depending on the conditions, whereas those carrying *nosZ* can also act as N_2_O sinks. Among incomplete denitrifiers, about 52 % had *nor* but not *nosZ*, whereas 34 % carried *nosZ*, either alone or in combination with *nir* or *nor* (53 and 28 % in the CXXCH clade, respectively; **Extended Data Table 2**). This suggests a higher N_2_O production than reduction capacity among *nrfA*-ammonifiers.

**Figure 2.**
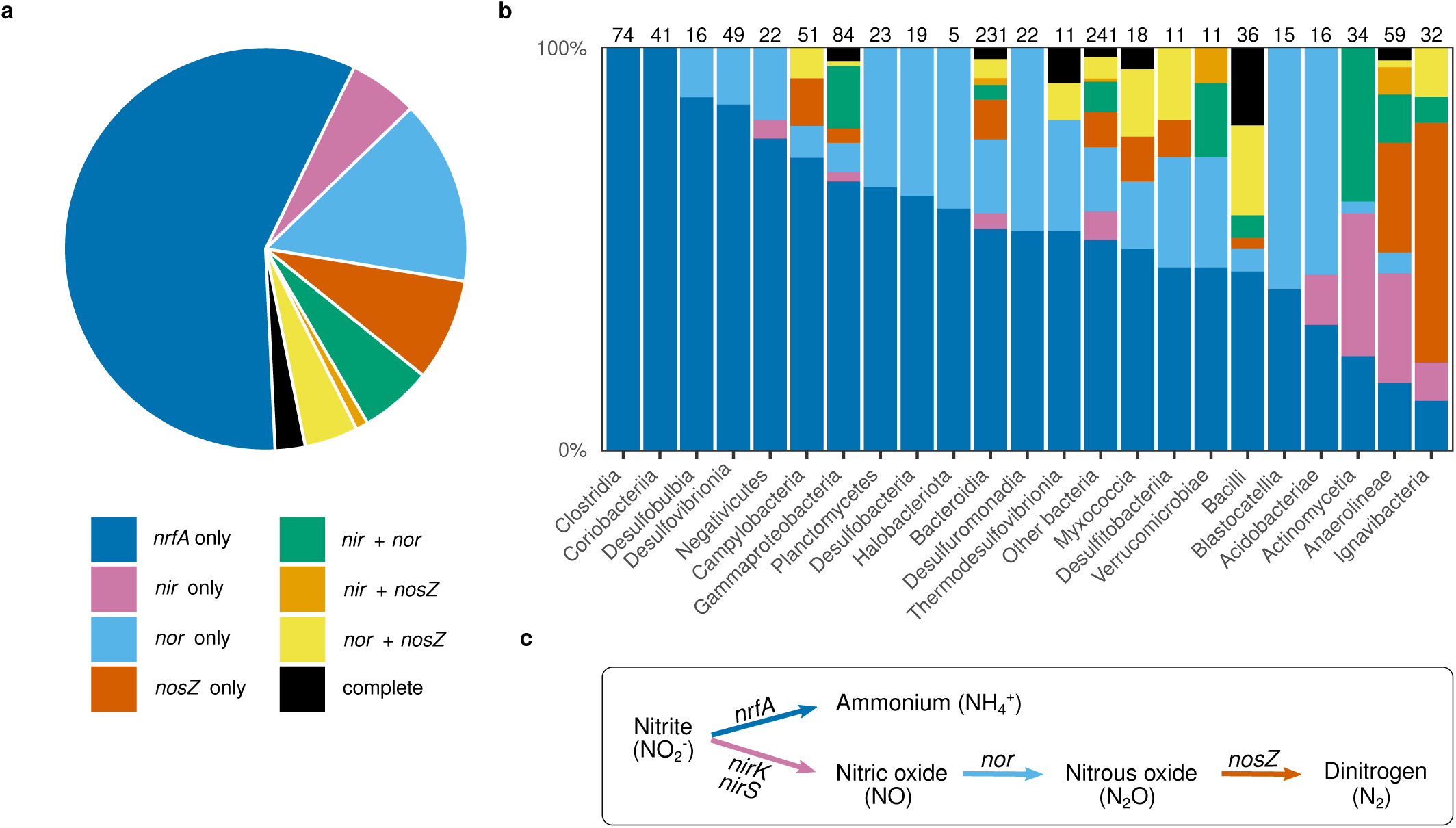
Co-existence of *nrfA* and denitrification genes in the 1,121 genome assemblies obtained when screening for *nrfA*. (a) Pie chart showing the distribution of *nrfA* and the denitrification genes *nirK*, *nirS*, *nor* and *nosZ* across all assemblies. (b) Distribution of *nrfA* and denitrification genes in the classes represented in the phylogeny in Fig. 1. The number of assemblies is indicated above the bar for each class. Classes are ordered according to the proportion of assemblies carrying only *nrfA*. (c) Reactions performed by the enzymes encoded by the different genes, with each arrow colored according to the corresponding gene.

Among the more frequently represented taxa in the *nrfA* phylogeny, the co-existence patterns between *nrfA* and denitrification genes displayed a large variation, ranging from 0 in Clostridia and Coriobacteriia to ca. 90% of assemblies carrying at least one denitrification gene in Anaerolineae and Ignavibacteria, and complete denitrifiers were predominantly found among Bacilli (**Fig. 2b**). Genomes with *nrfA* and genes coding for a nitric oxide (*nor*) or nitrite reductase (mainly *nirK*) were most common and evenly distributed across the phylogeny, whereas those with the nitrous oxide reductase gene (particularly *nosZ* clade II) was mainly restricted to Anaerolineae, Bacilli, Ignavibacteria and various members of the CXXCH clade (**Extended Data Fig. 2**).

### Environmental distribution and drivers of nrfA abundance

A collection of 1,861 globally distributed rhizosphere and soil metagenomes (**Fig. 3a**) was used to determine the abundance and diversity of *nrfA* communities across biomes. Reads corresponding to *nrfA* fragments were identified by mining each metagenome using a hidden Markov model of the reference alignment and candidate sequences were then mapped to the branches of the tree by phylogenetic placement to eliminate false positives corresponding to homologs (e.g. octaheme nitrite reductase fragments; Tikhonova *et al*., 2012). To account for differences in sequencing depth, *nrfA* placement counts were further normalized by the total number of base pairs sequenced in each metagenome (hereafter, ‘normalized *nrfA* counts’). The *nrfA* gene was present in all biomes, albeit in different proportions and in some cases with large within-biome variation (**Fig. 3b**). *nrfA*-ammonifiers were particularly prevalent in rhizosphere and croplands, with intermediate to low phylogenetic diversity (**Fig. 3b,c**). Tundra soils exhibited the highest phylogenetic diversity, but intermediate normalized *nrfA* counts. Forest ecosystems generally displayed a low abundance and a high diversity, except for tropical and subtropical dry broadleaf forest soils showing high normalized counts and low phylogenetic diversity. By contrast, normalized *nrfA* counts were lower in tropical compared to temperate and subtropical grasslands, savannas and shrublands, but with opposite patterns for phylogenetic diversity. The *nrfA* communities in desert soils were characterized by relatively low abundance and diversity. Overall, there was a negative correlation between normalized *nrfA* counts and phylogenetic diversity (*ρ* = -0.53; p < 0.001), mainly driven by rhizosphere and cropland communities (**Extended Data Fig. 3**).

**Figure 3.**
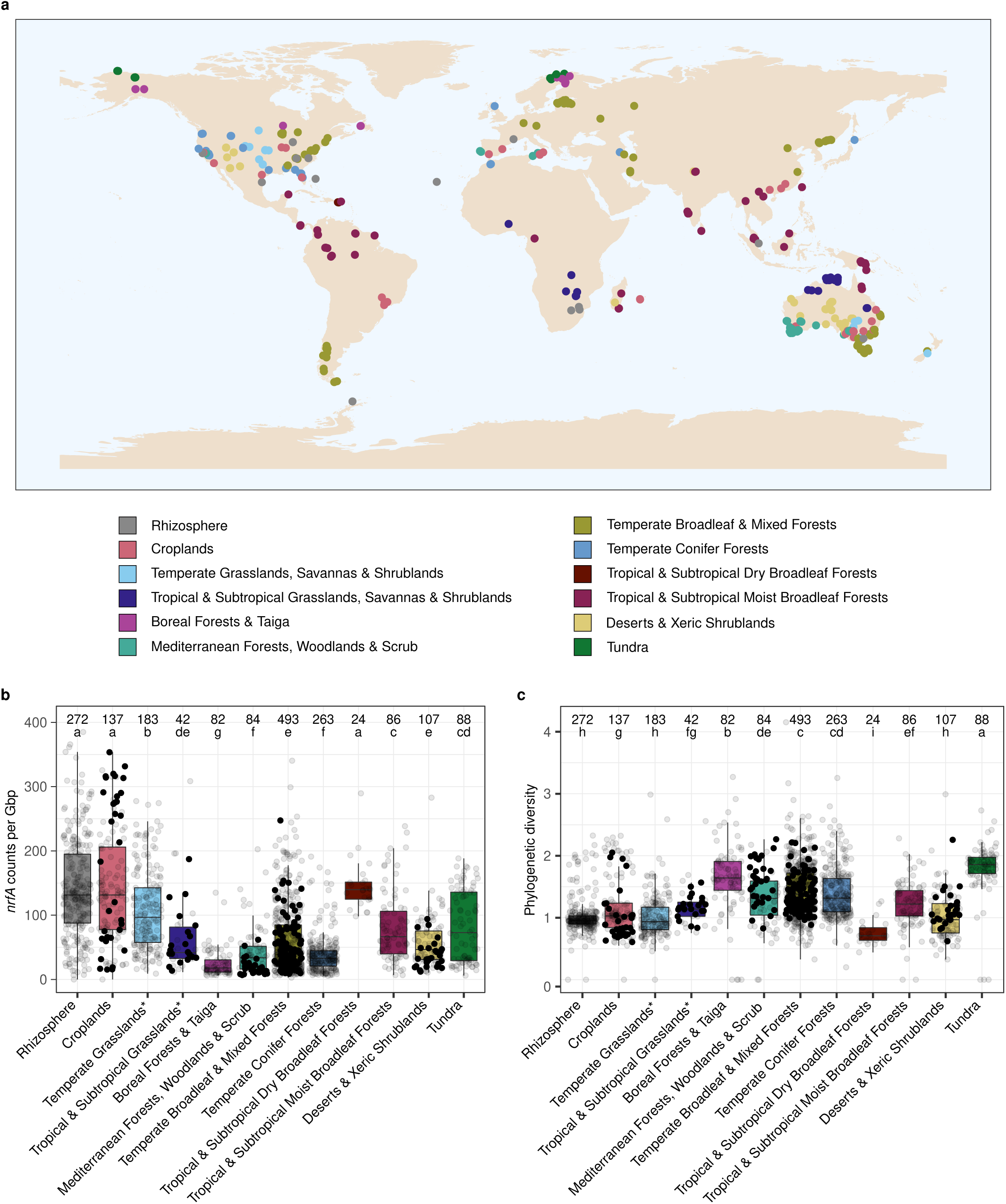
Location of metagenomes and abundance and phylogenetic diversity of *nrfA* across biomes. (a) 1,861 metagenomes representing 725 sampling sites across the globe. The 35 cropland and 5 rhizosphere metagenomes lacking associated geographic coordinates are not indicated. (b) Normalized *nrfA* counts per biome, calculated as the ratio between *nrfA* counts and the total number of base pairs (Gbp) sequenced in each metagenome (Kruskal-Wallis test, H = 641, P < 0.001). (c) Abundance-weighed phylogenetic diversity per biome (Kruskal-Wallis test, H = 576, P < 0.001). Significant differences are denoted with different letters, together with the number of metagenomes representing each biome above the boxplots. Boxes are bounded on the first and third quartiles; horizontal lines represent medians. Whiskers denote 1.5× the interquartile range. Data points corresponding to the metagenomes used in the random forest models are shown as filled circles. *The biome name also includes savannas and shrublands.

Phylogenetic placement of the *nrfA* sequence fragments on the reference tree showed that soil *nrfA* communities spanned the entire phylogeny but the CXXCH clade dominated across biomes (**Fig. 4a**). Most placements were located on deep branches in the phylogeny, indicating that abundant *nrfA*-carrying taxa in soil communities are distantly related to known *nrfA* representatives. Consequently, there was no biome-based discrimination of the *nrfA* communities in the phylogenetically-informed principal component analysis, and most of the variation in community composition was driven by different subsets of the CXXCH clade (**Fig. 4b**).

**Figure 4.**
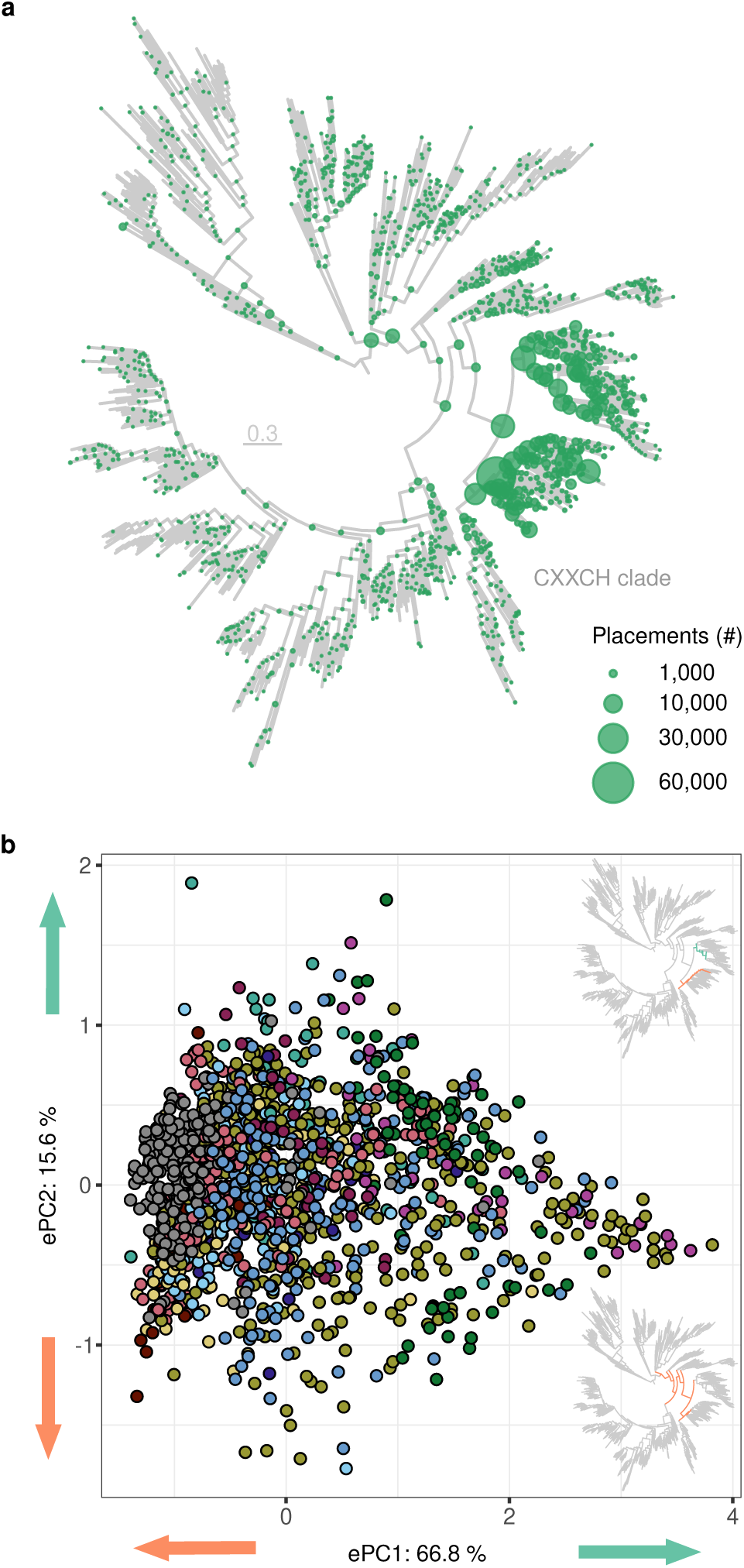
Phylogenetic placement of the metagenomic *nrfA* sequence fragments and associated phylogeny-based community composition across biomes. (a) Phylogenetic placement of the metagenomic *nrfA* sequence fragments on the reference tree. The size of the dots is proportional to the number of placements. The scale bar denotes the amino acid exchange rate (WAG+R10). (b) Edge principal component analysis showing differences in *nrfA* community composition between metagenomes grouped into biomes. Inset trees show which lineages are driving the separation of samples in positive (turquoise) and negative (orange) direction along each axis. The ordination was performed on metagenomes with ≥ 20 *nrfA* placements (n = 1,475). See Fig. 3a for the color legend of the biomes.

### Relative importance and drivers of NrfA-driven ammonification versus denitrification

The difference in normalized counts between *nrfA* placements and those of the marker genes *nirK* and *nirS* for denitrification was determined to assess the genetic potential for N retention at the community level in each of the 1,861 metagenomes (hereafter, ‘δ*nrfA*-*nir*’). Although the contribution of fungi to the genetic potential for denitrification was not included in the evaluation, a recent study has shown that denitrifying fungi have a minor influence on denitrification gene counts in terrestrial ecosystems (Bösch *et al*., 2022) and their omission is thus unlikely to affect the patterns reported here. All biomes exhibited a negative median δ*nrfA*-*nir* with few values above zero, indicating an overall lower genetic potential for NrfA-driven ammonification over denitrification (**Fig. 5a**). However, median values close to 0 were observed in both tundra and tropical & subtropical dry broadleaf forest soils, which in the latter appeared to be driven by high *nrfA* counts (**Fig. 3b**). Among forest biomes, the separation by climatic zones observed for the normalized *nrfA* counts (tropical > temperate ≥ Mediterranean > boreal) was not detectable in the δ*nrfA*-*nir* data, suggesting that conditions that favored *nrfA*-ammonifiers were even more favorable to denitrifiers. The median values of δ*nrfA*-*nir* in rhizosphere and cropland communities were at least 2-fold lower than in the other biomes. Among rhizosphere samples, tree (*Citrus* sp. and *Populus* sp.) and perennial grass (*Miscanthus* sp. and *Panicum virgatum*) species displayed higher δ*nrfA*-*nir* than the average, whereas bean (*Phaseolus vulgaris*) and *Brassica* spp. drove δ*nrfA*-*nir* towards higher abundance of denitrifiers. Maize (*Zea mays*) and thale cress (*Arabidopsis thaliana*) displayed δ*nrfA*-*nir* values comparable to the overall mean (**Fig. 5b**).

**Figure 5.**
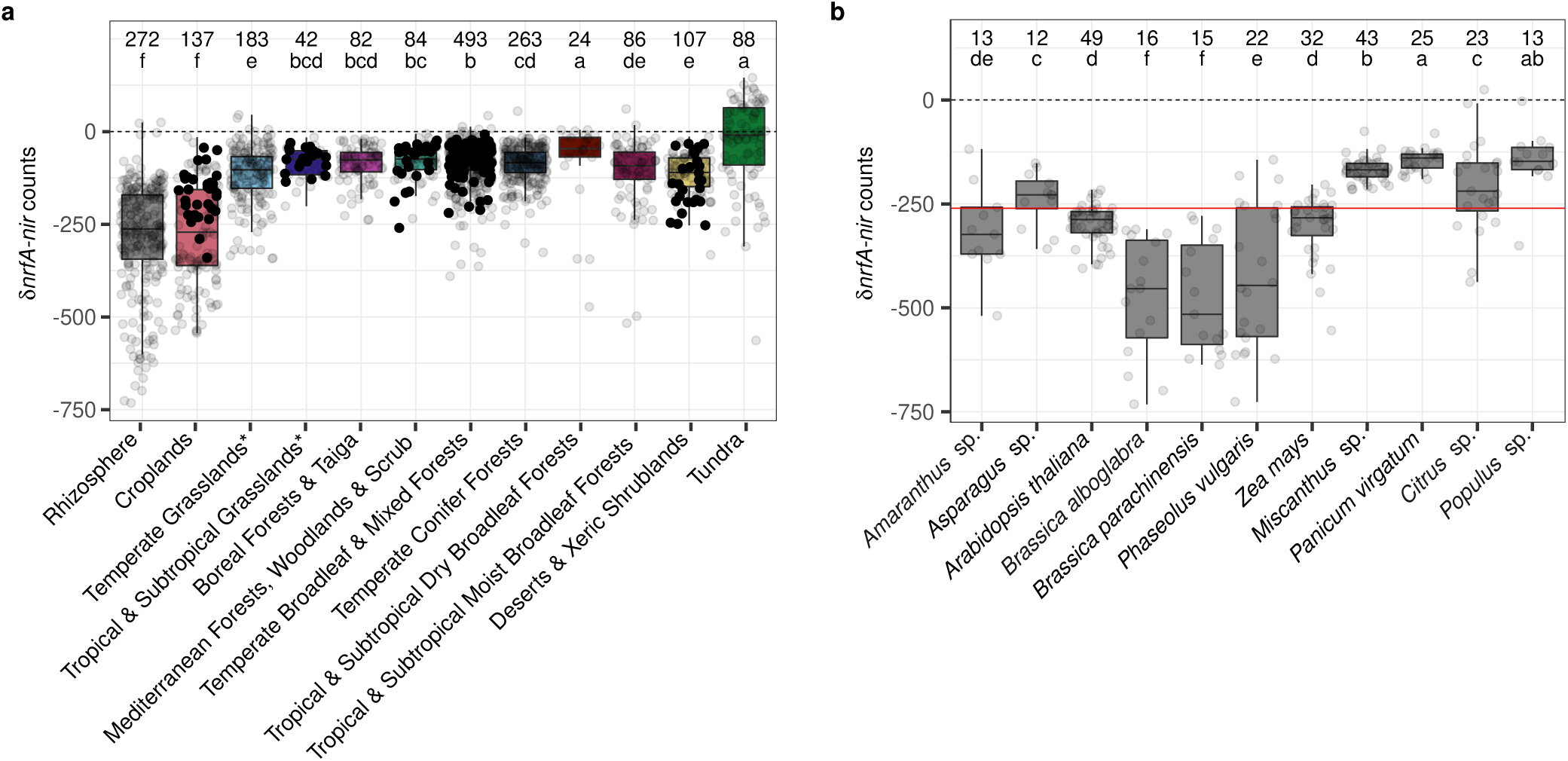
Relative importance of NrfA-driven ammonification and denitrification genetic potential across biomes. The difference in counts of *nrfA* and *nir* genes normalized by the number of base pairs sequenced (δ*nrfA*-*nir*) was calculated per metagenome. Positive and negative values indicate a higher potential for NrfA-driven ammonification over denitrification and vice versa. Significant differences are denoted with different letters, together with the number of metagenomes representing each biome. Boxes are bounded on the first and third quartiles; horizontal lines represent medians. Whiskers denote 1.5× the interquartile range. Data points corresponding to the metagenomes used in the random forest models are shown as filled circles. (a) Relative importance of NrfA-driven ammonification and denitrification genetic potential across terrestrial biomes and in the rhizosphere (Kruskal-Wallis test, H = 749, P < 0.001). (b) Relative importance of NrfA-driven ammonification and denitrification genetic potential in the rhizosphere of host species represented by more than 10 metagenomes (Kruskal-Wallis test, H = 174, P < 0.001). The red line indicates the median δ*nrfA*-*nir* value across all rhizosphere metagenomes. *The biome name also includes savannas and shrublands.

Environmental drivers of the normalized δ*nrfA*-*nir* counts were examined using random forest modelling on a subset of the metagenomes, which were selected based on the availability of metadata measured with the same methods across the metagenomes (Bissett *et al*., 2016) and including relevant factors for NrfA-driven ammonification and denitrification (**Table 1**). Accumulated local effect plots, showing the main effect of each variable in predicting δ*nrfA*-*nir* while controlling for other variables in the overall model, revealed a non-linear and overall positive relationship between the SOC to nitrate ratio and predicted δ*nrfA*-*nir*, mainly driven by low nitrate content rather than SOC levels (**Fig. 6**). Soil factors known to affect the activity of the different nitrite reductases also affected δ*nrfA*-*nir*. Calcium, essential for NrfA activity in the abundant CXXCH clade members (**Figs. 1** and **4a**) (Cunha *et al*., 2003), was associated with an increase in the prediction of δ*nrfA*-*nir* in the ranges 10-25 and 100-125 mM calcium kg^-1^, whereas copper, crucial for the activity of NirK (Zumft, 1997), had the opposite effect (**Fig. 6**). The δ*nrfA*-*nir* predictions were highest in acidic soils and displayed a u-shaped relationship with pH, indicating a threshold at 5.75 < pH < 6.5 and then increasing δ*nrfA*-*nir* with increasing pH in alkaline (pH > 7.5) soils, which aligns with measurements of nitrate ammonification rates across terrestrial ecosystems (Cheng *et al*., 2022). The genetic potential for NrfA-driven ammonification relative to denitrification was predicted to decrease with increasing phosphorus and sulfur (from 5 mg kg^-1^) concentrations. Biome identity was also identified as an important predictor, even after accounting for other environmental variables. Predictions obtained with the random forest models largely corresponded to the δ*nrfA*-*nir* calculated for the entire dataset, with predictions for croplands and deserts displaying lower relative potential for NrfA-driven ammonification compared to the average prediction, whereas forest and grassland soils had the opposite effect.

**Figure 6.**
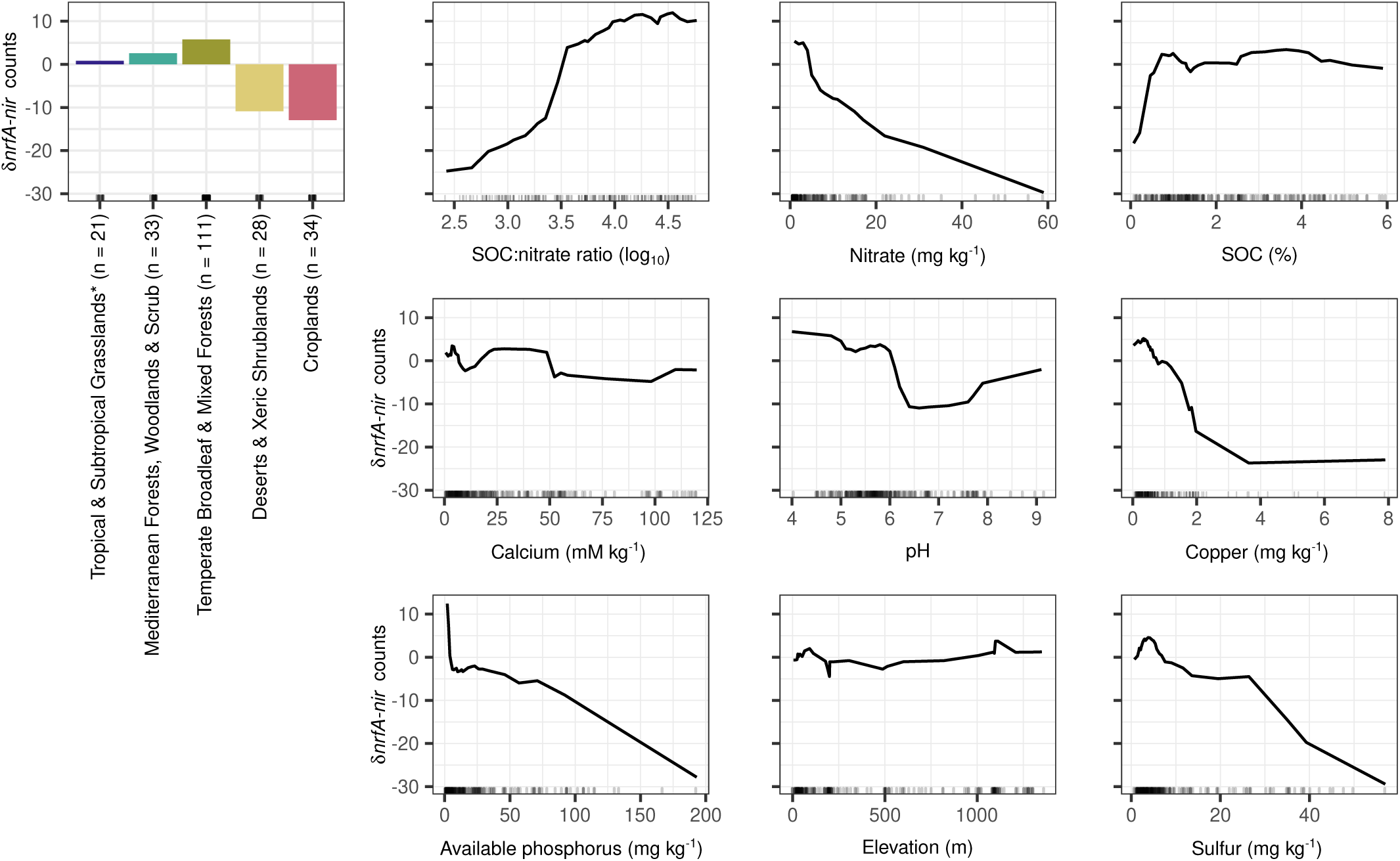
Environmental predictors of the potential competition between*nrfA*-ammonifiers and denitrifiers in soil based on random forest models. The difference in counts of *nrfA* and *nir* genes normalized by the number of base pairs sequenced (δ*nrfA*-*nir*) was calculated per metagenome. The analysis was performed on a subset of the metagenomes for which environmental metadata, especially soil properties relevant for nitrate ammonification and denitrification, was available (n = 227; Table 1). The number of metagenomes corresponding to each biome is indicated after the biome name. Predictor variables selected by VSURF and biome category were used to generate accumulated local effects plots, which show the differences in prediction of the δ*nrfA*-*nir* (y-axis) compared to the mean prediction along the range of each predictor (x-axis), while accounting for potential correlations amongst predictor values. The effect is centered so that the mean effect is zero. The random forest model was built with 500 trees, 2 features considered at each split and a tree depth set to 9 (variance explained: 55%, root mean square error: 40.6). SOC: soil organic carbon. *The biome name also includes savannas and shrublands.

**Table 1.** Continuous environmental variables associated to the “Biomes of Australian Soil Environments metagenomes”. (Bissett *et al*., 2016) **used in random forest modelling (n = 227).** Collinearity among soil and geomorphological factors was assessed by pair-wise Spearman correlations (**Extended Data Fig. 4**). The variable most relevant for nitrate ammonification and/or denitrification was retained in each collinear group (|r| ≥ 0.7). Variables used after the variable selection analysis are indicated in bold. The distributions of the gene counts and phylogenetic diversity data in the biomes covered by this dataset are shown by filled circles on **Figs. 3** and **5**.

| Category | Variable | Minimum | Maximum |
| --- | --- | --- | --- |
| Soil | Aluminium (mM kg<sup>-1</sup>) | 0.0 | 31.8 |
|  | **Ammonium (mg kg<sup>-1</sup>)** | 0.0 | 87.0 |
|  | Available potassium (mg kg<sup>-1</sup>) | 15.0 | 855.0 |
|  | **Available phosphorus (mg kg<sup>-1</sup>)** | 2.0 | 193.0 |
|  | **Calcium (mM kg<sup>-1</sup>)** | 0.6 | 158.1 |
|  | **Clay (%)** | 0.8 | 65.3 |
|  | Conductivity (dS m<sup>-1</sup>) | 0.0 | 8.9 |
|  | **Copper (mg kg<sup>-1</sup>)** | 0.0 | 32.7 |
|  | Iron (mg kg<sup>-1</sup>) | 1.9 | 1020.2 |
|  | Magnesium (mM kg<sup>-1</sup>) | 0.5 | 60.3 |
|  | **Moisture (%)** | 0.0 | 103.3 |
|  | **Nitrate (mg kg<sup>-1</sup>)** | 1.0 | 59.0 |
|  | **pH** | 4.0 | 9.6 |
|  | Silt (%) | 0.0 | 51.6 |
|  | Sodium (mg kg<sup>-1</sup>) | 2.3 | 6789.6 |
|  | **Organic carbon (%)** | 0.1 | 5.9 |
|  | **Sulfur (mg kg<sup>-1</sup>)** | 0.7 | 724.6 |
|  | Zinc (mg kg<sup>-1</sup>) | 0.0 | 32.4 |
| Geomorphology | **Elevation (m)** | 1.0 | 1350.0 |

## Discussion

Our gene-centric and phylogenetically-informed approach provides a framework for a more accurate understanding of the genetic potential for nitrate reduction in soil communities. We provide evidence that co-existence of *nrfA* and denitrification genes in genomes of *nrfA*-ammonifiers is common and phylogenetically widespread, with variation in gene combinations at fine phylogenetic levels. In contrast to earlier work suggesting that genetic capacity for N_2_O reduction is widespread among *nrfA*-ammonifiers (Sanford *et al*., 2012), our findings indicate that they are more likely to be N_2_O producers. This supports observations of N_2_O production from soil isolates of nitrate-ammonifying bacteria (Stremińska *et al*., 2012; Heo *et al*., 2020). The difference in genetic potential for N_2_O production *vs.* reduction was even more pronounced in the CXXCH clade, where most soil-derived *nrfA* reads were placed but not well represented among isolated and genome-sequenced *nrfA*-ammonifiers. Increased efforts aiming at characterizing members of the elusive CXXCH clade is needed to verify these patterns and understand other aspects of their ecology that are relevant to the cycling of N in terrestrial biomes. Key questions include whether differences in substrate affinity (Carlson *et al*., 2020; Wan *et al*., 2023) act as a major niche-differentiating factor in soil *nrfA*-ammonifiers, similarly to what is observed in other N transforming groups (Zhang & Okabe, 2020; Jung *et al*., 2022), and what role microbial interactions play in providing electron donors (e.g. low molecular-weight organic molecules by fermenters [Kraft *et al*., 2014] and nitrite for *nrfA*-ammonifiers lacking the ability to reduce nitrate [Wan *et al*., 2023]). Addressing these questions will both facilitate the interpretation of the links between measured rates of nitrate ammonification and *nrfA* community composition, diversity, and abundance, and help predicting how environmental factors influence the dynamics of the end-products when nitrate is used as an electron acceptor.

We show that the genetic potential for NrfA-driven ammonification was lower than that of denitrification overall in all terrestrial biomes, although the magnitude of the δ*nrfA*-*nir* counts varied both within and among biomes. Higher nitrate ammonification than denitrification rates have mainly been shown in sulfur-rich or reduced sediments (An & Gardner, 2002; Kessler *et al*., 2018; Murphy *et al*., 2020). However, we found similar patterns at the genetic level in the tundra sites, as many tundra metagenomes displayed positive δ*nrfA*-*nir* values. A high abundance of *nrfA*-ammonifiers has indeed been shown to support larger N stocks in these relatively N-limited ecosystems, which indicates that *nrfA*-ammonifiers play a role for N retention (Castaño *et al*., 2023). NrfA-driven ammonification could also be a relevant source of substrate for ammonia oxidizers, which potentially drive N_2_O emissions rather than denitrifiers in tundra soils (Ramm *et al*., 2022). Notably, managed systems (i.e. croplands and rhizosphere) exhibited significantly lower δ*nrfA*-*nir* than other soils and this difference was most likely due to fertilizer application (Putz *et al*., 2018), as nitrate and phosphorus were among the main drivers of decreased δ*nrfA*-*nir* in the random forest models. Moreover, lower δ*nrfA*-*nir* was observed in the rhizosphere of annual plants compared to perennials, which is consistent with the fact that cropping systems with perennial plants are typically characterized by comparatively more favorable conditions for nitrate ammonification, including higher SOC content (Ledo *et al*., 2020) and higher C/NO_3-_ratios, manifested in lower N_2_O production (Putz *et al*., 2018). The effect of SOC on predicted δ*nrfA*-*nir* rapidly reached a threshold (at ca. 0.8 % SOC), indicating a minor role for C quantity, but likely not C quality (Rütting *et al*., 2011; Carlson *et al*., 2020). Instead, the δ*nrfA*-*nir* was driven by low nitrate content, which aligns with *nrfA*-ammonifiers having a higher affinity than denitrifiers for nitrate or nitrite (van den Berg *et al*., 2016) and with the fact that C availability combined with low nitrate levels creates conditions suitable for nitrate ammonification, as more electrons are transferred per molecule of nitrate reduced (Strohm *et al*., 2007). Fertilized soils represent environments where the fate of N is most crucial to control (Steffen *et al*., 2015) and our findings suggest that integrating N management strategies promoting nitrate ammonification while also increasing soil carbon sequestration represents a promising way to increase N retention, especially in soils with low SOC content. This would reduce nitrate pollution and N_2_O emissions, while simultaneously increasing N use efficiency in agricultural soils.

## Online Methods

### Generation of reference nrfA and nir phylogenies

A previously published alignment of full-length amino acid NrfA sequences (n = 267; Cannon *et al*., 2019) was used to build a hidden Markov model (HMM) using the hmmbuild command implemented in HMMER v. 3.2 (Eddy, 2011). This model was used to screen the predicted ORFs in archaeal and bacterial genome assemblies available on RefSeq (NCBI, accessed in September 2019) for the presence of *nrfA* using hmmsearch (e-value cutoff = 1e-6). The resulting candidate *nrfA* sequences were translated to amino acids, dereplicated at 100 % identity using CD-HIT v. 4.8.1 (Fu *et al*., 2012) and aligned to the HMM model. The alignment was then manually inspected in ARB v. 7.0 (Ludwig *et al*., 2004) for the presence of conserved motifs representing catalytically important residues that are specific to the NrfA nitrite reductase (Simon, 2002). Due to the existence of related multi-heme cytochrome *c* proteins (Simon & Klotz, 2013), only full-length candidate NrfA sequences were retained at this step. The genome database was updated in October 2021 by downloading all archaeal (n = 8,131) and bacterial (n = 1,026,048) assemblies available on NCBI GenBank and screened using an updated HMM including the NrfA sequences identified in the 2019 search (n = 1,889). Candidate sequences were processed as described above and a set of octaheme nitrite reductase sequences retrieved by the final search was also retained to serve as the outgroup in the phylogeny (n = 84) (Soares *et al*., 2022). The alignment was refined by trimming the less conserved and poorly aligned C- and N-terminal regions and by removing columns with > 95 % gaps. FastTreeMP v. 2.1.11 (Price *et al*., 2010) was used to construct a draft phylogenetic tree and closely related sequences (i.e. very short terminal branch lengths) were manually pruned with the exception of *nrfA* copies originating from the same assembly. In the end, the curated data set contained 1,150 NrfA sequences originating from 1,121 assemblies. Selection of the best-fit model of evolution, WAG+R10, and construction of the amino acid-based maximum-likelihood phylogeny were performed with IQ-TREE v. 2.1.3 (Kalyaanamoorthy *et al*., 2017; Minh *et al*., 2020). Node support values were calculated using 1,000 ultra-fast bootstraps (Hoang *et al*., 2018) and the Shimodaira-Hasegaw approximate likelihood ratio (SH-aLRT) test (Guindon *et al*., 2010) with the -bnni option to reduce the risk of overestimating branch supports. The tree was plotted using iTOL v5 (Letunic & Bork, 2021). Reference phylogenies for NirK and NirS were generated using the same approach (**Supplementary Methods** and **Extended Data Figs. 5 and 6**).

### Quality check and taxonomic assignation of nrfA genome assemblies

The level of completeness and contamination of each of the 1,121 assemblies was determined based on the detection of lineage-specific, single-copy genes using BUSCO v. 5.2.2 (Manni *et al*., 2021). In the final dataset, 1,064 assemblies (ca. 95 %) met the high quality level standard (completeness > 90 % and contamination < 5 %) (Bowers *et al*., 2017). Lower quality assemblies corresponded to poorly sampled regions of the tree. Taxonomic annotations were obtained using GTDB-Tk v. 1.5.0 (Chaumeil *et al*., 2020) and the reference Genome Taxonomy DataBase r202 (Parks *et al*., 2020).

### Identification of denitrification genes in genome assemblies

The presence of denitrification genes (*nor*, *nirK*, *nirS* and *nosZ*) in the assemblies was also determined. HMM models for NirK, NirS and NosZ were built using alignments obtained by following the approach described for NrfA (**Supplementary Methods**), and for Nor the alignment from Murali et al. (Murali *et al*., 2022) was used. The models were used to search the final set of assemblies and the target sequences were identified by manually inspecting the resulting alignments.

### Construction of a metagenome database and GraftM search

A database was created by downloading 1,861 publicly available soil and rhizosphere metagenomes sequenced using Illumina short-read technology (read length ≥ 150 nt) and consisting of a minimum of 100,000 reads (**Extended Data Table 3** and **4**). The soil metagenomes were further classified into biomes, with the non-croplands (n = 1,462) classified following the definition of terrestrial ecoregions proposed by Olson et al. (Olson *et al*., 2001). Biome assignment was performed based on the geographic coordinates of each metagenome, using the ‘sp’ v. 1.4-6 (Pebesma & Bivand, 2005), ‘rgeos’ v.0.5-5 (Bivand & Rundel, 2021) and ‘rgdal’ v. 1.5-23 (Bivand *et al*., 2022) packages in R v. 4.2.0 (R Core Team, 2021). The geographic location of each sample was plotted using the ‘ggspatial’ v. 1.1.5 (Dunnington, 2021), ‘rnaturalearth’ v. 0.1.0 (South, 2017a), ‘rnaturalearthdata’ v. 0.1.0 (South, 2017b), ‘rgeos’ v.0.5-5 and ‘sf’ v. 1.0-7 (Pebesma, 2018) packages.

The presence of *nrfA* fragments in the metagenomes was assessed with GraftM v. 0.13.1 (Boyd *et al*., 2018), a gene-centric and phylogenetically-informed classifying tool. Using a custom gene reference package, GraftM identifies target gene fragments in metagenomes using HMMER and places them into a pre-constructed phylogenetic tree using PPLACER (Matsen *et al*., 2010). The phylogenetic placement acts as a validation step and ensures that the HMMER hits correspond to the target sequences and not to closely related outgroup sequences. A *nrfA* reference package for GraftM was built using the trimmed alignment and associated phylogeny generated in this study. However, sequences corresponding to genomic *nrfA* copies with very short terminal branch lengths were removed as they would increase computation time without increasing the sensitivity of the analyses (n = 1,104). Tree statistics required for running GraftM were calculated using RaxML v. 7.7.2 (Stamatakis, 2006) and the tree was re-rooted in iTOL v5 (Letunic & Bork, 2021).

The approach was first validated by shredding the aligned region of the *nrfA* sequences into 10,000 pieces of 150 nt-long fragments using GRINDER v. 0.5.4 (Angly *et al*., 2012). Full nucleotide octaheme nitrite reductase (outgroup) sequences were shredded in the same way. Each set of fragments was then individually processed with GraftM, using default parameters. GraftM provides up to 7 placements on the reference phylogeny for each read identified by HMMER and the ‘accumulate’ command implemented in gappa v. 0.8.1 (Czech *et al*., 2020) was used to find the most representative clade. Sensitivity and specificity were assessed by calculating the fractions of *nrfA* (ca. 87.5 %) and octaheme nitrite reductase (none) fragments placed into the ingroup, respectively. For comparison, 88.9 and 42.5 % of the *nrfA* and octaheme nitrite reductase fragments were identified as candidate sequences by the HMMER search, which stresses the importance of the placement step to eliminate false positives and shows the limited proportion of false negative generated by this approach. Sensitivity and specificity for *nirK* and *nirS* were 87.8 and 100 %, and 86.7 and 100%, respectively, with gene-specific outgroups for testing the specificity (various multi-copper oxidases for *nirK*; *nirN*, *nirF* and halophilic archaeal *nirS*-like sequences for *nirS*).

GraftM was run on the forward reads of each metagenome with default parameters using the first 150 nt only to account for differences in read length between metagenomes. Placements on the reference tree were visualized using the R package ‘ggtree’ v. 3.4.0 (Yu *et al*., 2017). In addition to *nrfA*, metagenomes were mined for denitrification (*nirK* and *nirS*) gene fragments. The resulting placement files were processed with gappa as described above. For each metagenome, *nrfA* counts were normalized by the number of base pairs (Gbp) sequenced to account for differences in sequencing depth (‘normalized *nrfA* counts’). The difference in normalized abundance (δ*nrfA*-*nir*) between *nrfA* and the marker genes for denitrification *nirK* and *nirS* was calculated as (*nrfA*-(*nirK*+*nirS*))/*Gbp*. The use of these metrics is relevant to assess the genetic potential for N retention in metagenomes since both *nrfA* (this study) and *nir* (Graf *et al*., 2014) genes are most commonly present in single copy in genomes.

### Statistical analyses

Abundance-weighed phylogenetic diversity (McCoy & Matsen, 2013), which provides a normalized measure of the shared phylogenetic history among taxa occurring in a sample, was calculated for each metagenome using the *nrfA* placements and the ‘fpd’ command implemented in the guppy suite of tools v. 1.1 (with θ = 1). The composition of the *nrfA* community across biomes was examined using the edge principal component analysis (Matsen & Evans, 2013) implemented in gappa (‘edgepca’ command). This ordination method is based on the phylogenetic placement of reads on a reference phylogeny and allows for the identification of specific lineages that contribute to the variation in composition between samples. It was performed on metagenomes with at least 20 *nrfA* placements as the algorithm otherwise failed to compute (n = 1,475).

All statistical analyses were performed in R v. 4.2.0. Differences in normalized phylogenetic diversity, *nrfA* counts and δ*nrfA*-*nir* across biomes were assessed using Kruskal-Wallis tests with multiple comparisons computed according to Fisher’s least significant difference and the false discovery rate correction available in the ‘agricolae’ package v. 1.3.5 (de Mendiburu, 2019). Figures were plotted using the ‘ggplot2’ package v. 3.3.5 (Wickham, 2016).

Relationships between environmental variables and δ*nrfA*-*nir* were determined using random forests, an ensemble machine learning algorithm that is well suited to model non-linear relationships between predictors and response variables and can deal with non-normality and high collinearity among predictors (Breiman, 2001). A subset of the metagenomes was selected based on the availability of soil metadata relevant to nitrate ammonification and denitrification, including pH and nitrate, organic carbon, calcium, and copper content (n = 227). All metagenomes within this subset belonged to the ‘Biomes of Australian Soil Environments’ project and covered large environmental gradients at the continental scale (**Table 1** and **Fig. 3a**). Since the corresponding samples were collected and processed following the same protocols (Bissett *et al*., 2016), this increased the likelihood to detect relevant ecological patterns. Collinearity among environmental factors was assessed by pairwise Spearman correlations (**Extended Data Fig. 4**) and only the most relevant variable for the processes in focus was retained in each collinear group (|r| ≥ 0.7; indicated in bold in **Table 1**). Random forest modelling was conducted as previously described (Saghaï *et al*., 2022). Briefly, data-driven variable selection was performed on the pre-selected environmental factors, supplemented with biome category information, using the ‘VSURF’ package v. 1.1.0 (Genuer *et al*., 2015) to identify the best predictors for δ*nrfA*-*nir*. The ‘randomForest’ package v. 4.7– 1 (Liaw & Wiener, 2002) was then used to model the relationship between the selected predictors and δ*nrfA*-*nir*. A grid search was first conducted to find the optimal combination of tuning parameters and the combination corresponding to the best model fit (lowest out-of-bag root-mean-square error) was selected. Results were then visualized using accumulated local effects plots (grid.size = 30) implemented in the ‘iml’ package v. 0.9.0 (Molnar *et al*., 2018; Apley & Zhu, 2020).

## Acknowledgments

The work was supported by a senior career grant to SH from the Swedish University of Agricultural Sciences and the Swedish Research Council Formas (grant 2019-00392 to SH). Part of the computational work and data storage was enabled by resources in projects NAISS 2021/23-527 and NAISS 2021/22-692 provided by the National Academic Infrastructure for Supercomputing in Sweden (NAISS) at UPPMAX, funded by the Swedish Research Council through grant agreement no. 2022-06725.

## Author contributions

SH conceived the study and acquired funding. AS, CMJ, GP and SH developed the methodological approach. AS, CMJ and GP built the phylogenies. GP performed the metagenome search. AS conducted the statistical analyses. AS and SH wrote the original draft and CMJ and GP provided critical feedback. All authors read and approved the final version of the manuscript.

## Data and materials availability

All data and code used in this study are present either in the main text, the Extended Data material or on Zenodo.

