## Extended Data File for "Phyloecology of *nrfA*-ammonifiers and their relative importance with denitrifiers in global terrestrial biomes"

**Extended Data Table 1.**

| Bacterial phylum | Number of assemblies | Archaeal phylum | Number of assemblies |
|---|---|---|---|
| AABM5-125-24 | 2 | Halobacteriota | 5 |
| Acidobacteriota | 51 | | |
| Actinobacteriota | 78 | | |
| Armatimonadota | 11 | | |
| Bacteroidota | 287 | | |
| Bdellovibrionota | 9 | | |
| Calditrichota | 1 | | |
| Campylobacterota | 51 | | |
| Chloroflexota | 61 | | |
| Chrysiogenetota | 2 | | |
| CLD3 | 2 | | |
| Cyanobacteria | 2 | | |
| Deinococcota | 7 | | |
| Desulfobacterota | 125 | | |
| Elusimicrobiota | 4 | | |
| Eremiobacterota | 1 | | |
| FEN-1099 | 1 | | |
| Fibrobacterota | 1 | | |
| Firmicutes | 182 | | |
| Gemmatimonadota | 6 | | |
| Goldbacteria | 1 | | |
| Hydrogenedentota | 6 | | |
| Krumholzibacteriota | 3 | | |
| KSB1 | 3 | | |
| Latescibacterota | 1 | | |
| Marinisomatota | 1 | | |
| Methylomirabilota | 2 | | |
| Myxococcota | 37 | | |
| Nitrospinota | 2 | | |
| Nitrospirota | 15 | | |
| OLB16 | 1 | | |
| Omnitrophota | 1 | | |
| Planctomycetota | 31 | | |
| Poribacteria | 1 | | |
| Proteobacteria | 88 | | |
| QNDG01 | 1 | | |
| Spirochaetota | 12 | | |
| Sumerlaeota | 4 | | |
| UBA10199 | 3 | | |
| UBP17 | 1 | | |
| UBP4 | 1 | | |
| Verrucomicrobiota | 15 | | |
| Zixibacteria | 2 | | |

**Extended Data Table 2.**

| | Gene or gene combination | Number of assemblies |
| --- | --- | --- |
| 444 incomplete denitrifiers in 1,121 assemblies | *nir* only | 62 |
| | *nor* only | 166 |
| | *nosZ* only | 92 |
| | *nir* + *nor* | 65 |
| | *nir* + *nosZ* | 11 |
| | *nor* + *nosZ* | 48 |
| 177 incomplete denitrifiers in 309 assemblies in the CXXCH clade | *nir* only | 31 |
| | *nor* only | 59 |
| | *nosZ* only | 27 |
| | *nir* + *nor* | 36 |
| | *nir* + *nosZ* | 7 |
| | *nor* + *nosZ* | 17 |

**Extended Data Table 3.**

| Biome | Number of metagenomes | Reference |
|---|---|---|
| Croplands | 41 | Bisset et al. GigaScience 2016 |
| | 25 | Hartman et al. ISME J 2017 |
| | 3 | Mendes et al. ISMEJ 2017 |
| | 12 | Orellana et al. Appl Env Mic 2018 |
| | 36 | PRJNA717057 |
| | 20 | Xu et al. Nat Com 2020 |
| Deserts and Xeric Shrublands | 3 | Bahram et al. Nature 2018 |
| | 39 | Bisset et al. GigaScience 2016 |
| | 65 | NEON* |
| Boreal Forests & Taiga | 21 | Wilhem et al. Sci Data 2017 |
| | 7 | Bahram et al. Nature 2018 |
| | 54 | NEON* |
| Mediterranean Forests Woodlands and Scrub | 17 | Bahram et al. Nature 2018 |
| | 45 | Bisset et al. GigaScience 2016 |
| | 22 | NEON* |
| Temperate Broadleaf and Mixed Forests | 21 | Wilhem et al. Sci Data 2017 |
| | 66 | Bahram et al. Nature 2018 |
| | 139 | Bisset et al. GigaScience 2016 |
| | 244 | NEON* |
| | 12 | Sorensen et al. Nat Microbiol 2019 |
| | 11 | PRJNA717057 |
| Temperate Conifer Forests | 44 | Wilhem et al. Sci Data 2017 |
| | 8 | Bahram et al. Nature 2018 |
| | 59 | Diamond et al. Nat Mic 2019 |
| | 152 | NEON* |
| Tropical and Subtropical Dry Broadleaf Forests | 24 | NEON* |
| Tropical and Subtropical Moist Broadleaf Forests | 78 | Bahram et al. Nature 2018 |
| | 6 | Bisset et al. GigaScience 2016 |
| | 2 | Mendes et al. ISMEJ 2017 |
| Temperate Grasslands Savannas and Shrublands | 2 | Bahram et al. Nature 2018 |
| | 12 | Bisset et al. GigaScience 2016 |
| | 169 | NEON* |
| Tropical and Subtropical Grasslands Savannas and Shrublands | 13 | Bahram et al. Nature 2018 |
| | 29 | Bisset et al. GigaScience 2016 |
| Tundra | 5 | Bahram et al. Nature 2018 |
| | 27 | NEON* |
| | 56 | Woodcroft et al. Nature 2018 |

*NEON (National Ecological Observatory Network). Soil microbe metagenome sequences (DP1.10107.001), RELEASE-2021. https://doi.org/10.48443/fzzj-g053.

**Extended Data Table 4.**

| Plant host species | Number of metagenomes | Reference |
|---|---|---|
| *Amaranthus* sp. | 13 | Bandla et al. Scientific Data 2020 |
| *Arabidopsis thaliana* | 49 | Levy et al. 2017 Nature Genetics |
| *Asparagus* sp. | 12 | Crovadore et al. Genome Announc 2017 |
| *Phaseolus vulgaris* | 22 | Mendes et al. ISME J 2017 |
| *Brassica alboglabra* | 16 | Bandla et al. Scientific Data 2020 |
| *Brassica parachinensis* | 15 | Bandla et al. Scientific Data 2020 |
| *Citrus* sp. | 23 | Xu et al. Nat Com 2018 |
| *Colobanthus quitensis* | 3 | Molina-Montenegro Polar Biology 2020 |
| *Colobanthus quitensis* + *Deschampsia antarctica* | 3 | Molina-Montenegro Polar Biology 2020 |
| *Zea mays* | 32 | PRJNA330341-47, PRJNA367156-68, PRJNA405457, PRJNA406023-27, PRJNA444376-80 |
| *Gossypium* sp. | 1 | Singh et al. Microbiol Resourc Announc 2020 |
| *Miscanthus* sp. | 43 | PRJNA330359-60, PRJNA365493-99, PRJNA366147-53, PRJNA366178-79, PRJNA367152 -53, PRJNA375575-80, PRJNA405458-61, PRJNA444381-85 |
| *Populus* sp. | 13 | Blair et al. mSystems 2018 |
| *Helianthus annuus* | 1 | Babalola et al Data in Brief 2020 |
| *Panicum virgatum* | 25 | PRJNA330352-58, PRJNA365487-92, PRJNA375569-74, PRJNA405463-67, PRJNA444386 |
| *Taxus cuspidata* | 1 | Hao et al. J. Basic Microbiol 2018 |

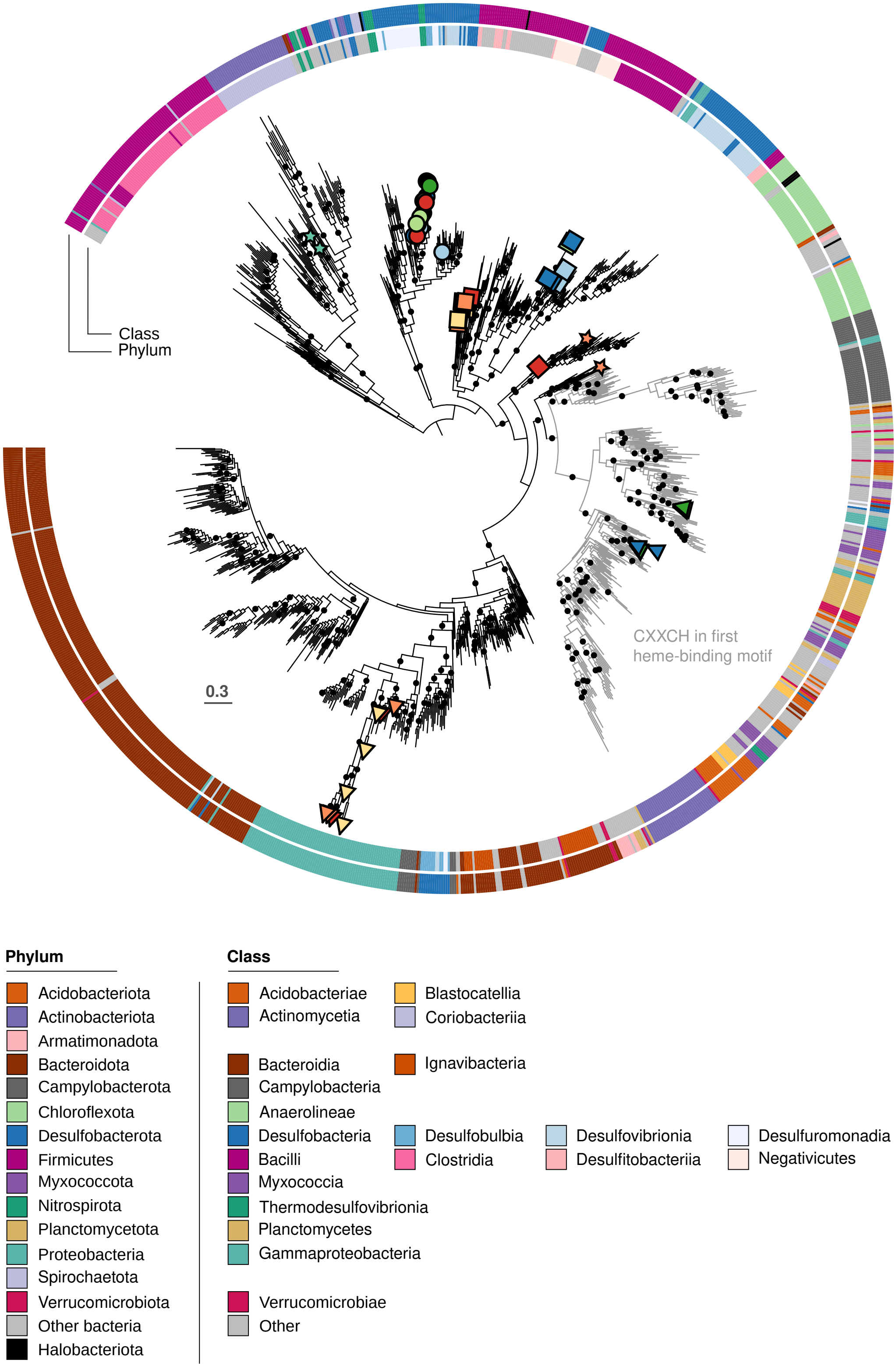


**Extended Data Figure 1.**

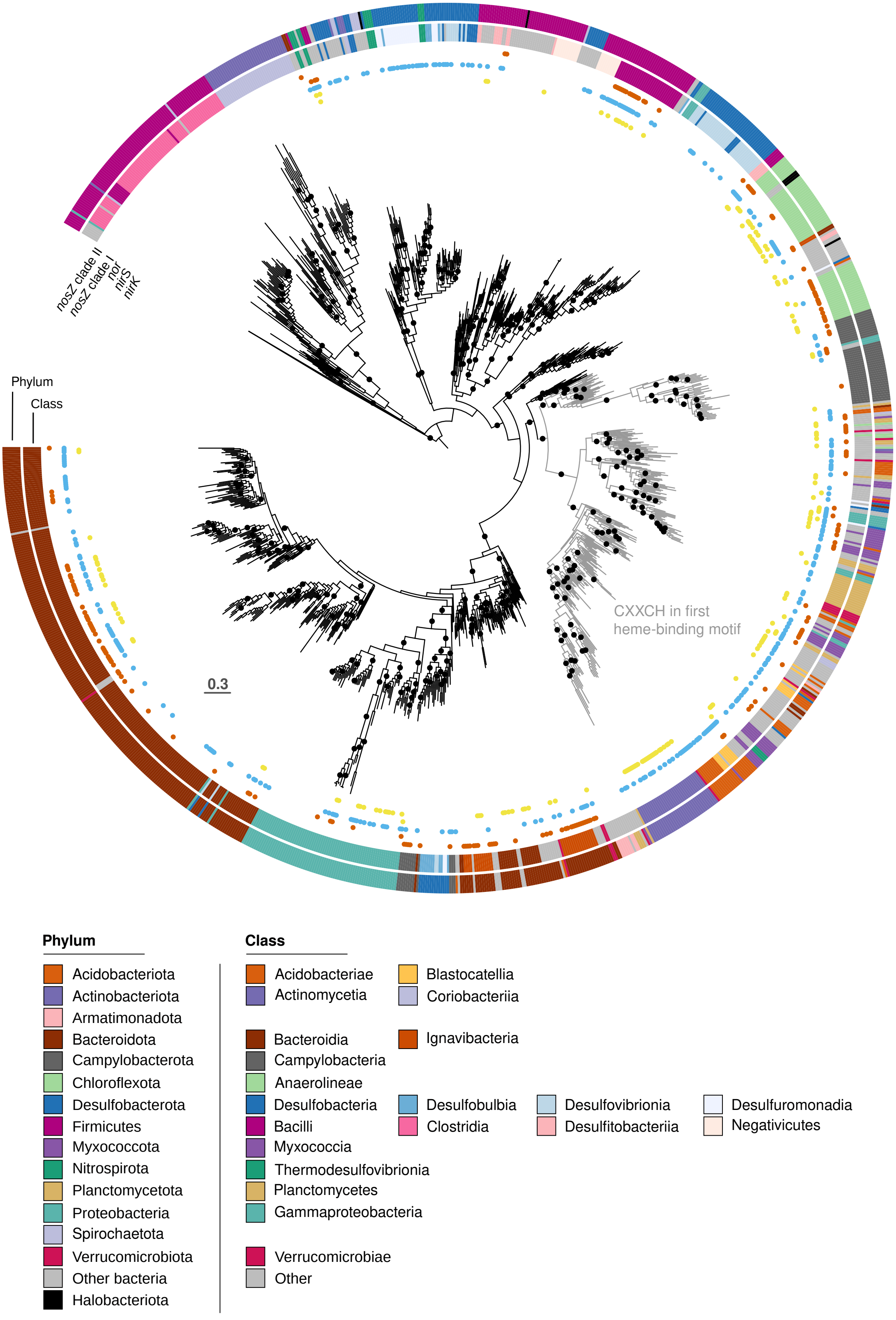


**Extended Data Figure 2.**

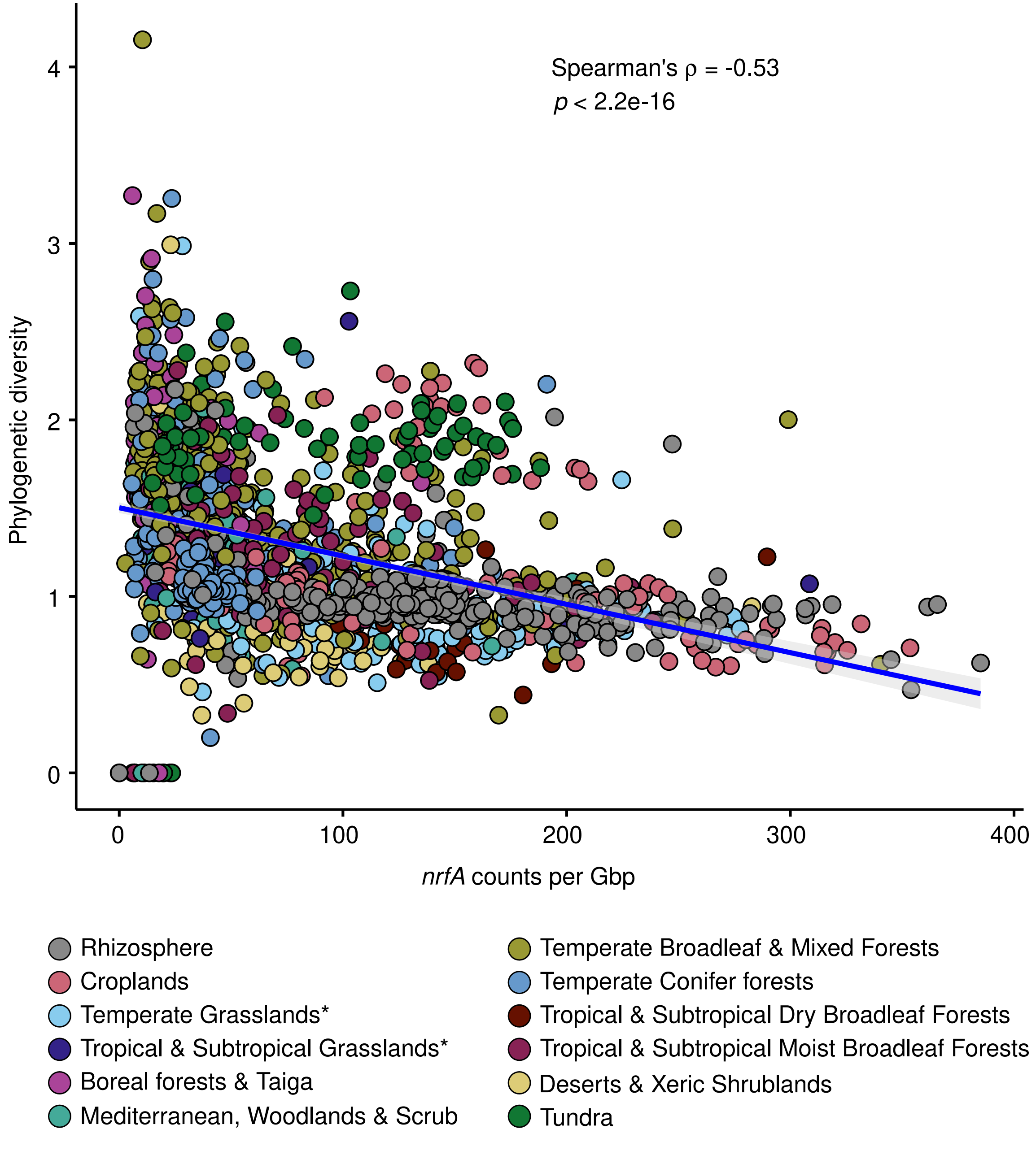


**Extended Data Figure 3.**

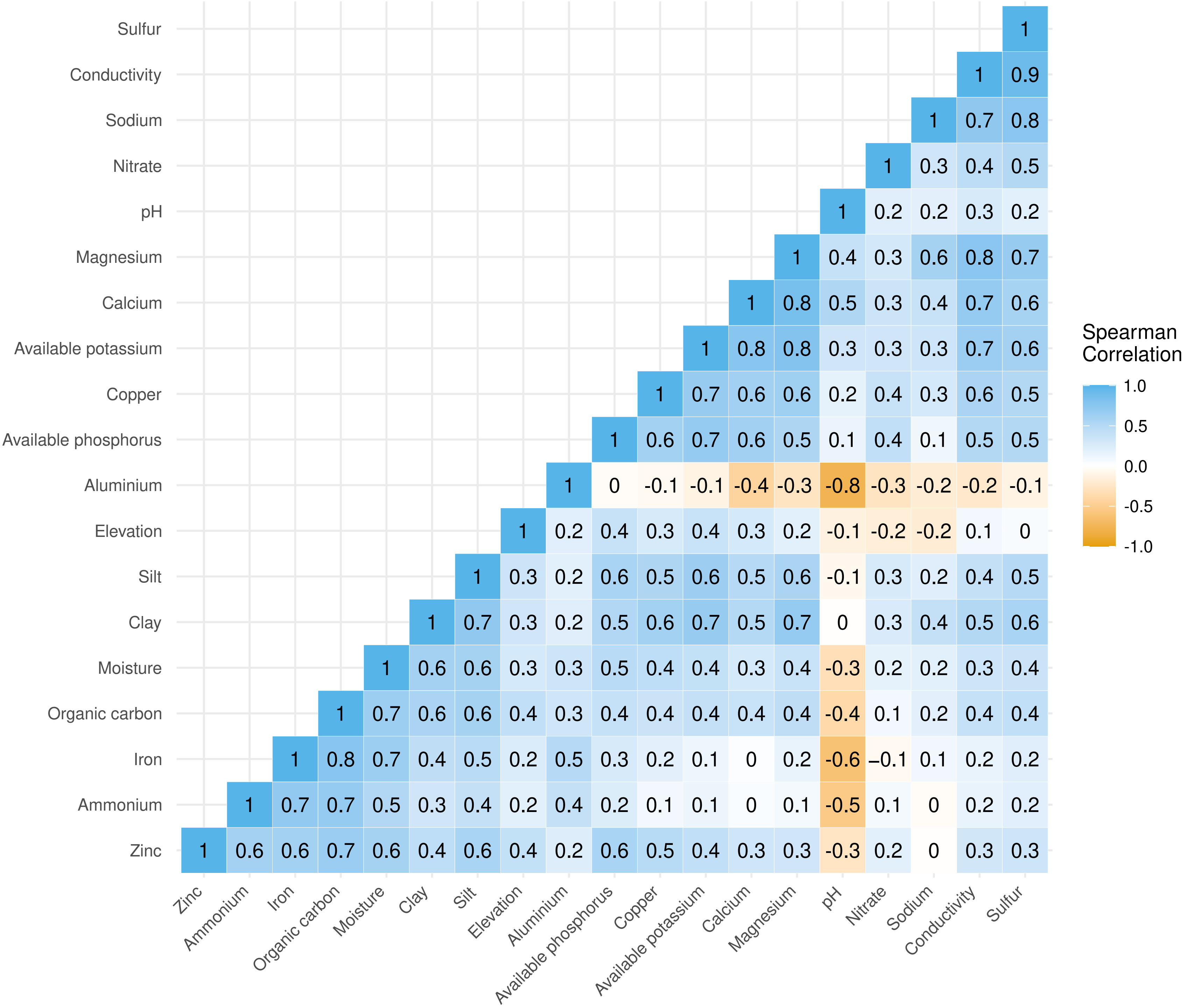


**Extended Data Figure 4.**

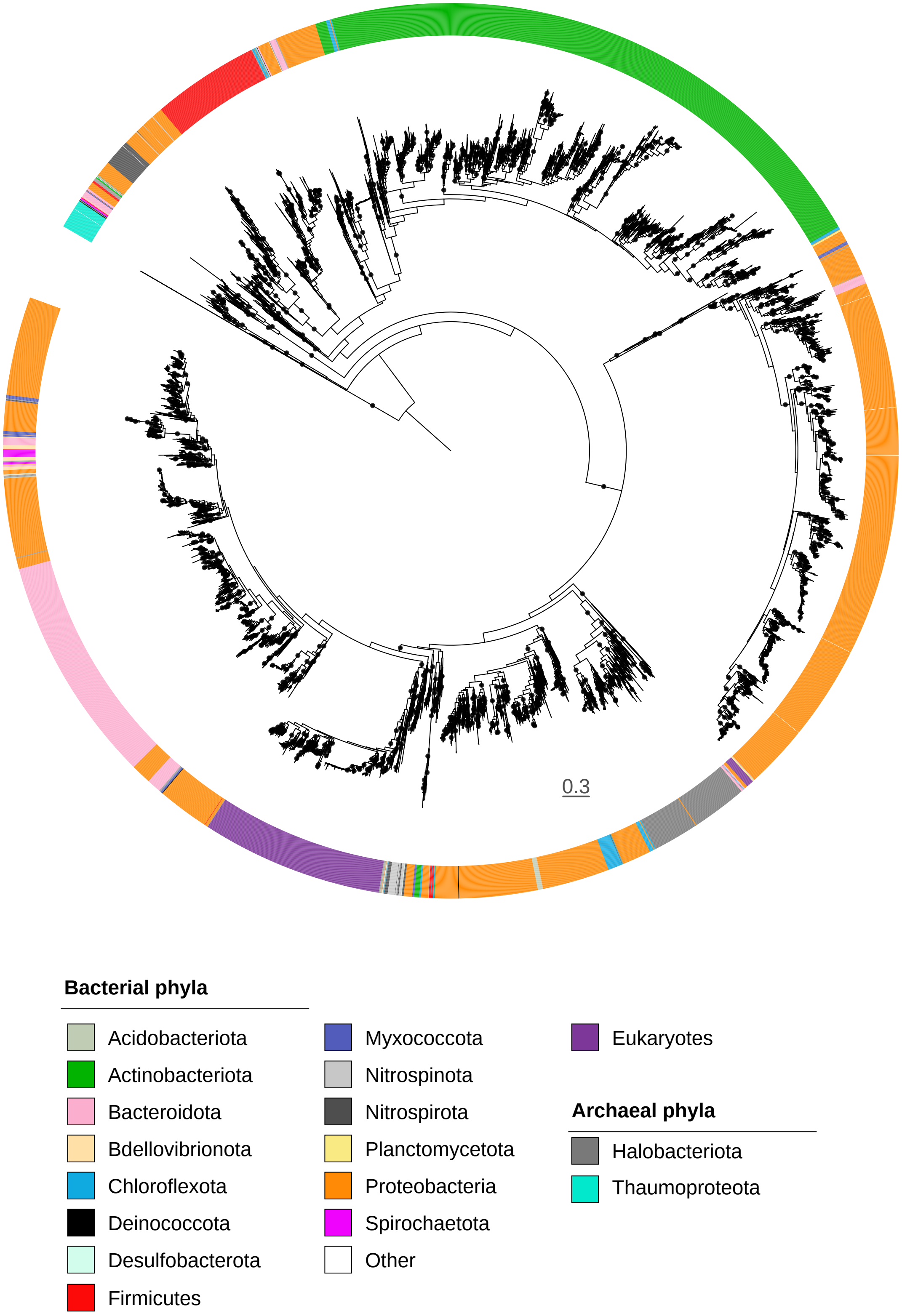


**Extended Data Figure 5.**

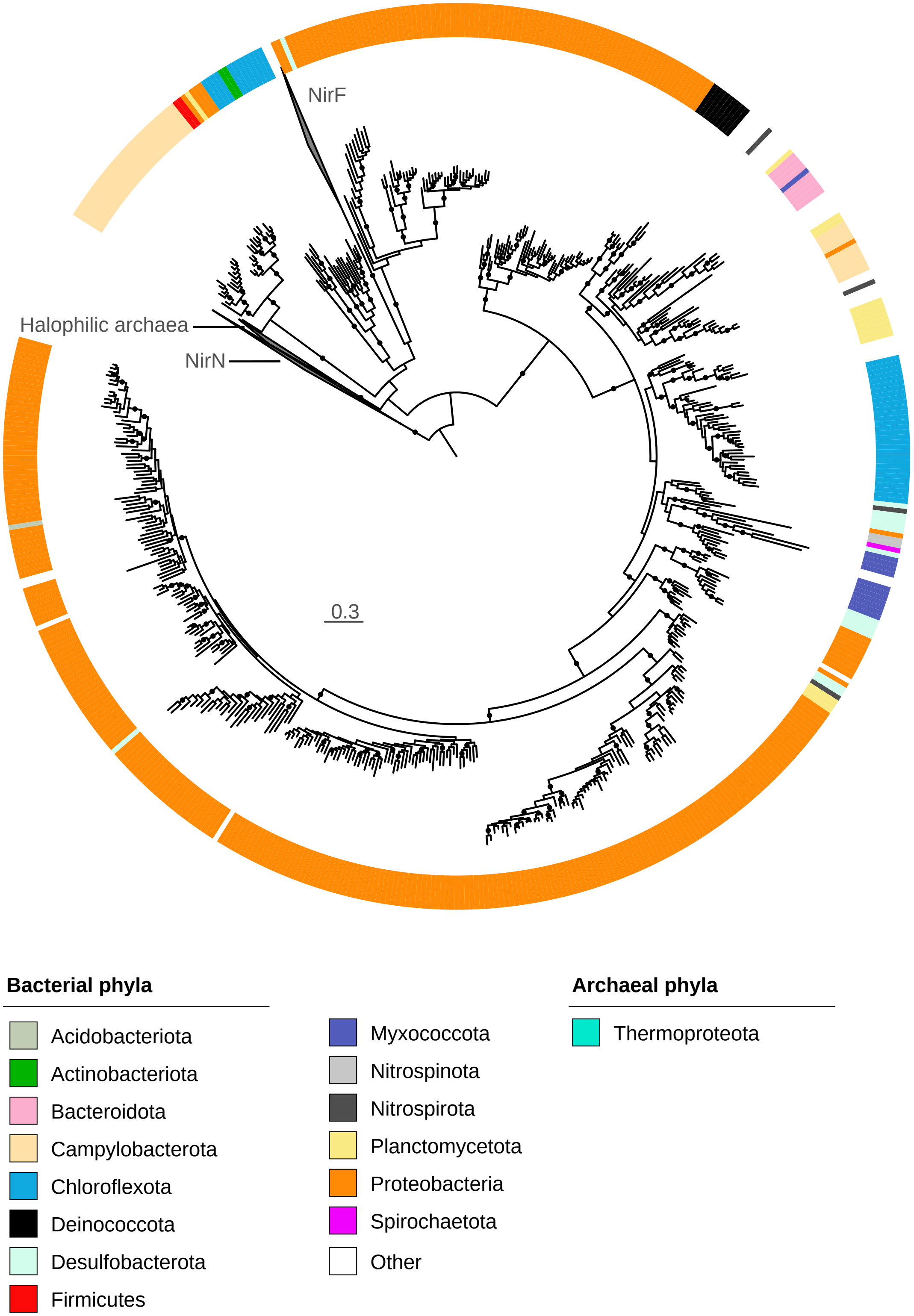


**Extended Data Figure 6.**
